# Cell Centred Finite Element Model for Intestinal Organoids Shape Analysis: From tissue architecture to mechanics

**DOI:** 10.1101/2022.01.27.477994

**Authors:** J. Laussu, D. Michel, S. Segonds, S. Marguet, F. Barreau, D. Hamel, E. Mas, A. Ferrand, F. Bugarin

**Affiliations:** Institut Clément Ader,Université Fédérale de Toulouse Midi-Pyrénées, Institut Clément Ader – CNRS UMR 5312 – UPS/INSA/Mines Albi/ISAE, 3, rue Caroline Aigle, Toulouse 31400, France; IRSD, Université Fédérale de Toulouse, INSERM, INRAE, ENVT, UPS, U1220, CHU Purpan, CS60039, 31024 Toulouse, France; Centre de réeférence des maladies rares digestives et service de Gastroentérologie, Hépatologie, Nutrition, Diabétologie et Maladies Héréditaires du Métabolisme, Hôpital des Enfants, CHU de Toulouse, F-31300, France

**Keywords:** Cell Shape, Biomechanics, Cell Mechanics, Vertex Model, Organoid, FEM

## Abstract

Organoids, established from stem cells owing to their self-renewal and differentiation capacities, are self-organised three-dimensional tissue culture recapitulating the original cell populations and their associated functions, as well as tissue architecture. The intestinal organoids established from adult stem cells isolated from the intestinal crypts, recreate 3D epithelial mini-intestines. They represent an excellent tool to study intestinal stem cell capacities and their ability to reconstitute a fully polarised and functional epithelium. These 3D cultures recapitulate *in vitro* the tissue characteristics, including architecture, either in physiological or disease conditions whether they are established from healthy or pathological tissue samples respectively. In this regard, their use for potential treatments screening carries the hopes for a future personalized medicine for which image-analysis such as HCS are increasingly being developed. Numerous numerical models have been developed to study the effects of organoid development on their shape. Most of them remain mainly restricted in their physical description due to the complex inter-relationship between cell physics, phenotypes and behaviors, exploding the number of variables in modeling formulation. Finite Element Method (FEM) is a numerical analysis method employed in mechanics to model deformation and evaluate residual stress of complex structures making it difficult to obtain analytical solutions. Considering epithelial architecture as a homogeneous material where each cell is an elemental equivalent part of the problem, FEM allows a direct link between tissue architecture deformations and local mechanical constraints. Here we formalise a new organoid cell centred FEM with a physical description borrowed from the engineering world. This model can allow a better understanding of the individual contribution of physical/mechanical properties of individual cells on general tissue architecture.

**Author Summary:** Stem cell research has experienced a breakthrough with the development of powerful tools, namely, organoids. The possibility to recreate *in vitro* from adult stem cells a fully functional adult tissue with its actual 3D architecture made accessible poorly defined steps in tissue morphogenesis or pathophysiology. If recent efforts have focused on studies of tissue specific organoids functions, modelling a generic bio-mechanic model of organoid remains necessary to decipher the mechanical biodiversity of organoid structures.

## I. INTRODUCTION

Understanding the interplay between biology and physics in epithelium architecture for highly renewal tissue is a challenging question. The intestinal epithelium undergoes continuous self-renewal and requires a delicate balance to keep the epithelial barrier integrity. The experimental efforts to understand this homeostasis have largely focused on the study of the regulation of cell proliferation, cell differentiation and cell death balance [1, 2]. However other parameters like cell shape regulation [3, 4], cell migration [5] or osmotic pressure [6] have recently been proved involved in epithelial tissue shaping. Deciphering the individual contributions of these highly intertwined functions necessitates to produce a powerful computational model bringing together biology, physics and geometry. This requires also to choose an appropriate biological support, allowing multi-scale observations from cellular resolution to global tissue functions and architecture.

Nowadays exist some powerful biological models designed to question stem cell capacities to establish a fully organised and functional tissue, the organoids [7]. Organoids are autonomous and self organised systems that recapitulate the 3D original tissue architecture establishment, including epithelium polarisation or folding for instance [8]. Their abilities to integrate complex cell behaviours in a relatively short timeline make these *ex vivo* mini-organs perfect tools to study how biology and physics of the cell cooperate to establish functional tissues, or how tissue architecture alteration can be linked to diseases initiation and progression.

Although theoretical efforts on tissue morphology have largely been driven by 2D experiments, recent models are interested in 3D processes occurring in complex morphogenetic events [9–11]. Among them, spatial models, dealing with volumes, shapes and mechanics of the cells, have shown themselves relevant for organoid systems modeling [12], a large majority being dedicated to digestive organoids [13]. The development of these computational models, focusing on organoids or simple sheets as a “model of a model”, represent a perfect combination between computational prediction and rapid experimental validation. In various 3D vertex epithelial models, cells are generally represented as a polyhedron, a boundary face between neighboring cells expressed by a polygon. The tissue, at a macroscopic level, is a cell aggregate expressed by a single edges network. For dynamical modelling (4D models), this network can be iteratively rearranged in order to report on tissue deformation relative to the changes in cell topology during proliferation. These vertex models include 3D and 2D vertex models [14]. In 2D, surface polygon meshes only represent the surface as polygons, when in 3D, volumetric meshes discretise the interior volume of a cell. This model can be used for sub-cellular entities if polygons are smaller than the cell. Using a bigger polygon, an over-cellular model gives access to tissue level description. This general use of Vertices and Vertex to construct the computational model allows to distinct three major categories depending on their structure: 1/ Structured Grid models are lattice-based and space is divided up into a discrete grid, in most cases a regular lattice. These continuous models, with no inter-spaces, are ideal to study self-organisation of the tissue with the diffusion of chemical gradients [15]. However, the lack of real cell shape information makes it inappropriate to model physics and cell shape relationship at cellular level. 2/ Unstructured Grid models and use of Off-lattice grids have huge advantages for morphogenesis studies. Cell shape is not constrained to a predefined regular shape (e.g. hexagonal in most Drosophila models) and can dynamically evolve in the model. Moreover, theses continuous models have proven a useful formalisation of mechanical behavior for tissue cells packing or confluence [14, 16–18]. 3/ Finally No Grid models or Agent Based models are symmetrically opposed in their approach of the modelling. They use discrete tissue organisation, decomposing the general problem in a sum of individual problems. The general base of these models are spheres representing cells and allowing the elaboration of dynamical theories over time, like fluidity in cell migration, cell differentiation or growth rate [19, 20]. These models are powerful for tissue patterning but the discontinuity in the geometrical structure makes it difficult to use for mechanical properties studies [21]. Considering this, we choose to use an unstructured Grid Model with volumetric meshes to model our bio-mechanical organoid model.

The use of continuous vertex models has a huge advantage for mechanical structure analysis with one particular application of volumetric meshes, Finite Element Analysis (FEA) [22]. FEA uses regular or irregular volumetric meshes to compute internal stresses and forces throughout the entire volume of an object. The Finite Element Method (FEM) is a well-established numerical analysis method employed in mechanics to model deformation of complex structures in which it might be difficult to obtain an analytical solution. FEM is well designed to model the effect of deformation due to local constraints. If we reverse the problem we can link local constraint to deformations and this reverse problem is ideal to infer forces or mechanical properties of epithelial tissue, which are rarely accessible with classical invasive methods by the morphological survey of the tissue. FEM uses large collections of predefined elements to approximate by polynomial functions on each element, the complete partition. To sum up, FEM proposes a discrete solution of a continuous problem. FEM was rapidly employed for solid tissue modelling [23]. Regarding epithelial architecture modelisation, tissue can be considered as a continuous material wherein each cell is an elemental equivalent part of this problem. Here, the global shape of an organoid can be divided in geometrically simple elements inside the cells to ensure both a sub-cellular mechanical record and a good approximation of the cell shape. The cells remain the principal entities of the system integrating all the cellular components associated to specific behaviour, nucleus, cell membrane, cytoplasm, cellular cortex, using as many elements needed.

Morphology is one of the principal features defining intestinal organoids [24]. Here we propose to use the volumetric vertex model of organoids associated with FEM in order to simulate the organoid architecture alteration using simple physical law and material properties. We present the conception and the formulation of our vertex model organised at a sub-cellular level and subsequently we validate our organoid modeling using an existing active vertex model. Finally, we demonstrate how flexible the model is and may provide a basis for linking cell shape, tissue deformation and stress at a cellular level due to imposed constraints, allowing the addition of mechanical information in accordance with previous vertex epithelial tissue models.

## II. RESULTS

### A. Biological Reference

We based our model on intestinal organoids deformation. Colon tissues are obtained from patient endoscopic biopsies. After isolation, the crypts are cultured in matrigel matrix support during 24 hours to obtain colon organoids (Fig. 1A.a). Colonic organoids come from a nearly spherical organisation, during cystic step, with a poor number of large cells, to a complex building organoid following cell compaction due to their polarisation (Fig. 1B). The polarisation is not achieved for all the organoids at the same moment, some of them remaining cystic. If we can measure a cell apical and basal surface only defined by the surface at the respectively external and lumen interface, these two terms ‘apical’ and ‘basal’ are biologically relevant only after cell polarisation events visible with actin accumulation at the apical surface [25](Fig. 1C.a-b). We can clearly see the evolution of the global shape of the organoid correlating with restructuration of individual cell shape with three distinct steps, each being associated to a specific basal-lateral cell aspect ratio (Fig. 1C.a). The first one entitles ‘cystic organisation’ where basal-lateral cell aspect ratio is above 1, a second phase of transition corresponding in the polarisation of the tissue around a ratio of 1. And a third step of ‘columnar organisation’ with compact polarised cells, associated with a loss of the radial symmetry and the appearance of some evagination (Fig. 1C.b). All these observations show the highly dynamic remodelling capacities of the organoid model relative to it’s simple architecture and illustrate the necessity to use multi-scale models for tissue deformation (macro scale) requiring precise cell shape information (micro scale). This also drives the choice to use the geometrical cell property ‘basal-lateral cell aspect ratio’ first as a global parameter to design distinctly cyst or columnar organoids, and second as a potential local curvature (deformation) information for the rest of our approach.

**Figure 1:**
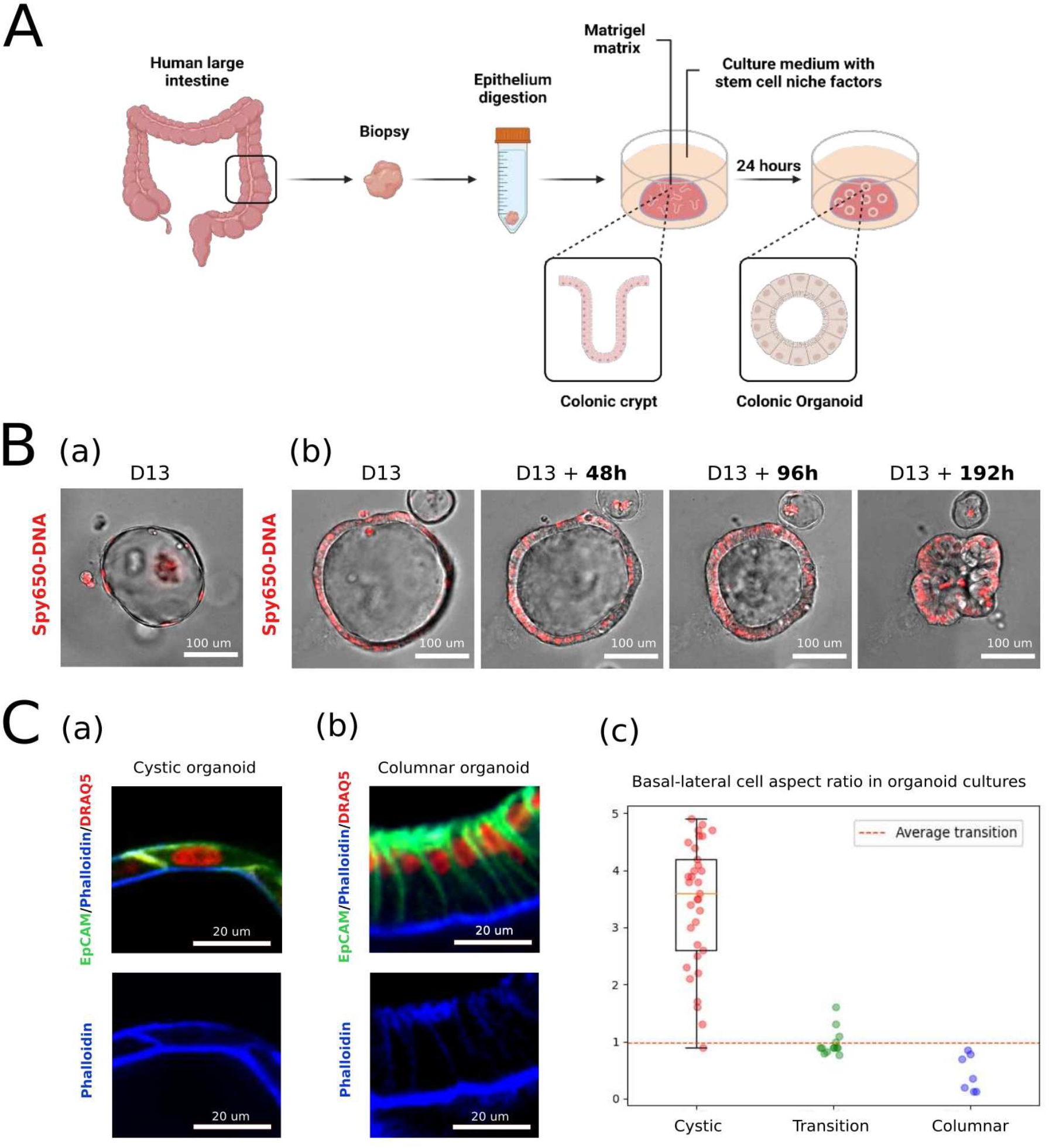
Colonic organoids present a dynamical architecture involving cell shape changes. A/ Schema representing the experimental process to obtain colonic organoid from Human biopsies after isolation and culture in matrigel matrix of the crypts. Illustration was created with BioRender.com. B/ Growth pattern of human colonic organoids in Matrigel on a timelapse analysis with widefield and nucleus live label Spy650-DNA (red): (a) cystic conformation of an organoid acquired 13 days before polarisation from a spherical cystic organisation to a collapse phase resulting in the polarisation of cells, (b) after polarisation from a spherical columnar organisation to an other collapse phase prelude of the budding expansion. C/ High magnification at cellular scale of a snapshot imaged on a fixed columnar organoid with phalloidin (blue) immunostaining, EpCAM (green) immunostaining, and nuclear Draq5 (red) staining (a) a ‘cystic organoid’ after 15 days of culture in matrigel or (b) a columnar organoid after 15 days of culture. (c) Distribution chart of the lenght/height cell aspect ratio observe from the following of 10 organoids during 240 hours time lapse experiment. A plateau around equal lenght/height (=1) cell aspect ratio corresponds to the polarisation phase. Above organoids are majority cystic and bellow they are columnar.

**TABLE I:**
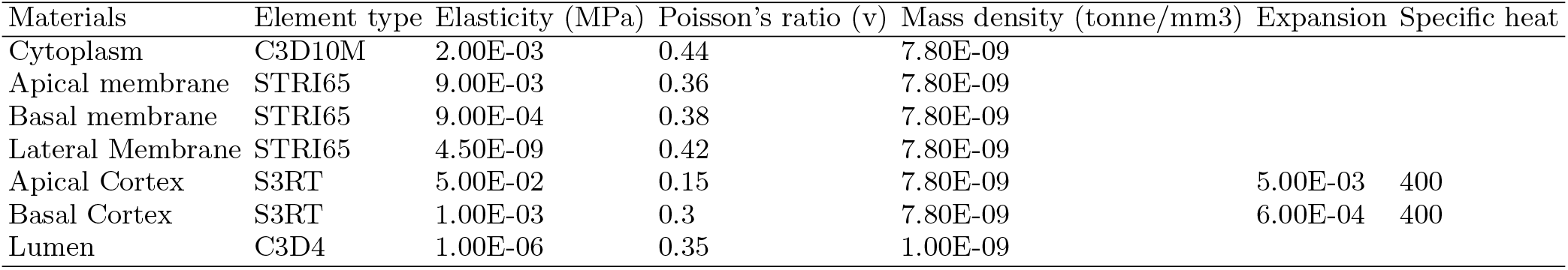
FEM Properties

### B. Mesh generation

#### a. Mesh generation

In our model, the cell is a polygon mesh, a collection of vertices, edges and faces. Faces are the external structures of the cells, both cell-cell and cell-exterior interfaces. Edges are the segments at the perimeter of the polygonal faces. Finally vertices are the ensemble of edge nodes. All these parts are dependent to a cell identity and can be addressed from vertices to cells or recursively without losing any part. To construct virtual organoids, and because the initial default structure of organoids is spherical, we decide to distribute cell positions uniformly using fibonacci sphere distribution. The sphere size was normalized and only the cell positions number figure as parameters. After that we introduce three different steps of meshing to obtain the cell volumes. First we performed a Delaunay tessellation to connect neighboring seeds followed by a Voronoi tessellation and delimit cell area in 2.5D (Fig. 2A) to create a simple shell. Finally we duplicate and connect two copies of this flat shell to create a volumic closed tissue monolayer. Thickness of the monolayer is defined as a ratio of the radius of the principal sphere; this parameter, 0.3 x radius, was arbitrarily fixed at the creation of the virtual organoid in order to confere an basal-lateral cell aspect ratio ¡1 to model columnar organoids (Fig. 1C.a). This new virtual epithelial sheet was polarised defining as apical the faces exposed to internal lumen of the spheroid and basal the faces exposed to external environment, which is the Matrigel matrix in organoid culture conditions. The meshes were exported to a FEM for Abaqus©. Abaqus© is a commercial finite element solution software that was used to obtain the deformation or strain produced in the process of organoid mechanical FEA.

**Figure 2:**
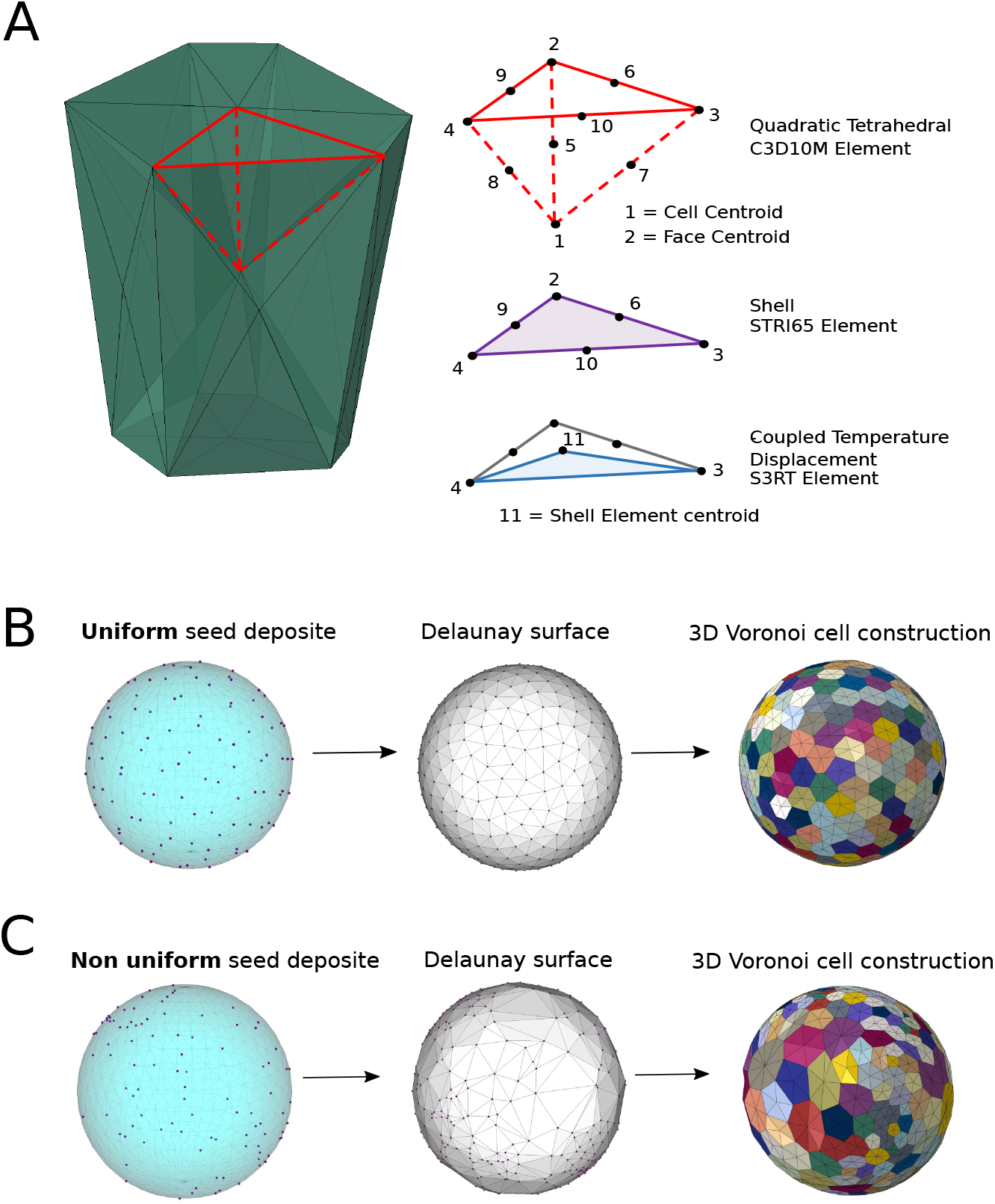
Construction of the virtual organoid for Finite Element Analysis. A/ Description of the subcellular element model. Cell are composed of three different types of elements: Quadratic tetrahedral elements for the cytoplasm volume discretisation using C3D10M predefined Abaqus© element, quadratic Shell elements for the membrane discretization using STRI65 predefined Abaqus© element and coupled Temperature-Displacement triangles for the cortex discretization using S3RT predefined Abaqus© element only for the apical and basal surface of the cells. B/ Uniforme virtual organoid are generated following from three successive steps: (a) seeds are evenly distributed using the mapping of a Fibonacci lattice on a sphere (b) a Delaunay 3D creating convex hull of the surface (c) Voronoi 3D on 2 spherics convex hull with 2 distinct radius to create a volume. All cells have more or less the same volume. C/ Non-uniforme virtual organoid are generated following from three successive steps: (a) seeds are randomly distributed on a sphere (b) a Delaunay 3D creating convex hull of the surface (c) Voronoi 3D on 2 spherics convex hull with 2 distinct radius. Cell volumes variation is important.

#### b. Finite element formulation

We constructed individual cells as a collection of three types of interactive subcellular elements without any shape *a priori* (e.g. basal shape can be triangular, hexagonal, or a more complex polygon independently to the apical shape). In that perspective, we choose tetrahedral elements, more convenient than hexahedral elements to cover uniformely a sphere [26] and more flexible for complex cell shapes [27](Fig. 2A). - Cytoskeleton: cells intra part are segmented with quadratic C3D10M Abaqus© tetrahedral element connecting the centroid of the cell and a part of the face of the cell (Fig. 2A). - Membranes: external cells interfaces are modeled with quadratic STRI65 Abaqus© shell elements connecting only the nodes of the preceding volumic elements present at the surface of the cells (Fig. 2A). - Apical and Basal Cortex: acto-myosin network are modeled for apical and basal surface only, with first order S3RT Abaqus© thermal shell elements (Fig. 2A). All these elements are suffisant to create complex cell shape thanks to the use of quadratic elements allowing more degree of freedom and the implementation of all specific properties used in common vertex models (e.g. volumic elasticity, surface elasticity, surface tension). Finally the inner part of the organoid is a fluid-filled cavity meshed in 2 layers with first order tetrahedral elements (C3D4 Abaqus© element) in order to confer volume elasticity to the lumen.

#### c. Model parameters

One of the originality of our model, arising from use of FEM, is to confer material properties to the living tissue. Here, we defined the physical behavior of the FEM organoid in four distinct domains, cell cytoplasm, cell membrane, cell cortex, and the extracellular lumen, each using specific elements, e.i. specific mechanical properties (I). For simplicity, it is assumed that cell cytoplasms are purely elastic [28] and this elasticity is isotropic. In the literature, epithelial cell cytoplasm elasticity is characterized with a Young’s modulus ranging from 0.3 to 100 kPa depending on the biological model and the measurement method [20, 28–30]. For cytoplasm elasticity in all our models, we fixed an arbitrary value of 2 kPa, after validation of simple sheet deformation, associated with a Poisson’s ratio of 0.44. Indeed, high Poisson’s ratios (near 0.5) are largely used for deformations which occur at constant volume. Similarly, we gave an elastic behavior to the membrane elements in order to represent membrane elasticity based on the literature [30, 31]. Due to the different nature of the interface, we give a higher elasticity to the apical membrane compare to the basal membrane which is surrounded by a basal lamina. The last surface of the cell, lateral membrane, at the interface between neighboring cells, has the particularity to connect two cellular membranes. In our continuous vertex model, we do not duplicate membranes for cell-cell contacts in order to simplify the problem. With this unique interface, we reduced the number of contacts to attribute and compute during solving iterative operations. The constraints at this lateral membrane interface are neglected using a low Young’s modulus of 4.5E-06 kPa. By this way, the constraints at cell-cell interfaces are mostly due to pressure equilibrium exercise by cytoplasm volumes.

Concerning the cytoskeleton, we defined apical and basal triangles to represent respectively apical and basal actin-myosin cortex. In a lot of tissues, and this is the case for colonic organoids after polarisation of the epithelium (Fig. 1), cortical actin-myosin network is enriched at the apical surface of the cell compared to the lateral-basal surface. We decided to make this distinction by creating two different surfaces, apical cortex and basal cortex and give them two distinct coefficients of expansion representing the relative acto-myosin concentration in our model. With two times higher elasticity for basal cortex and near ten times higher expansion coefficient for apical cortex we tend to mimic this relative distribution of actin activity. Finally, to model lumen incompressibility, lumen volume elasticity was decomposed in a very low Young’s modulus and a relatively high Poisson’s ratio representing respectively the deformability and the incompressibility of the medium. All these values are given as an order of value and because we are focusing our study on the geometrical and mechanical relationship relevance, we are using virtual homogeneous organoids (all mechanical properties are similar from a cell to the other one).

#### d. Active cell model

We assumed that cytoplasm acts like a nearly incompressible fluid with a nearly undiminished diffusion of the stress inside the cells, this is why changes in cell shape can be summed up by contraction or relaxation of their cytoskeleton [32]. Complex cell architecture is subjected to dynamical transformation over time as a mechanical response to environment changes. That is even more true when cells need to divide or move but also to maintain tissue homeostasis against stress. Apical constriction is an important cell shape change that promotes tissue remodeling in a variety of homeostatic and developmental contexts (e.g. gastrulation and neural tube formation in vertebrates) [33]. In recent years, progress has been made towards understanding how the distinct cell biological processes that together drive apical constriction are coordinated. This process of apical contraction requires for a part the contraction of actin-myosin network itself [33], which generates force, and in addition the cell-cell junctions, which allows forces to be transmitted between cells. Because apical actin-myosin enrichment and contractility have become defining characteristics of apical constriction mechanisms in which epithelial cells undergo apical shrinkage [34], we decide to implement this function in our 3D epithelial FEM model by the use of a thermal formulation. When a solid material is heated, it expands depending on two principal properties, a specific heat and a rate of expansion. This rate of expansion is not linear and the result for thermal stress increases according to a fixed coefficient of thermal expansion. In our model, we decide to use this thermal property to model cell contractility. Given a lower temperature to thermal elements, we aim to contract them to mimic actin contraction or, on the contrary, relaxation in heating the cortex elements. With this term, thermal induction and rate of expansion can be respectively representative of the pulse of contraction and the concentration of actin-myosin protein present at the cortex.

**TABLE II:**
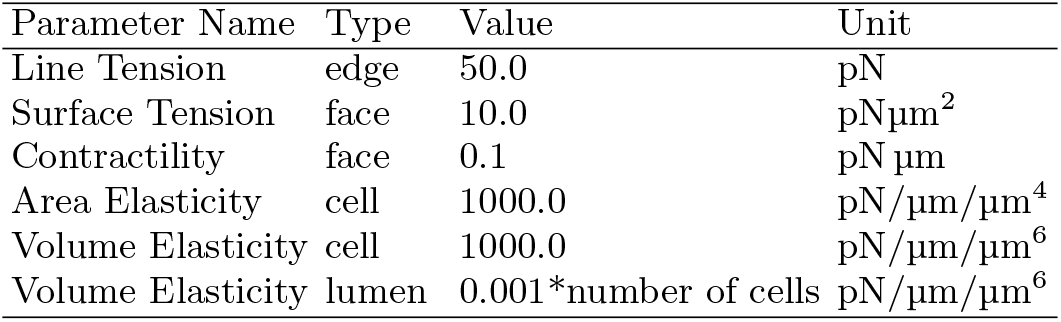
Rheology Model parameters

### C. Static model for shell shape analysis

Morphogenesis and tissue renewal are essentially driven by differential growth resulting from cell divisions and cell rearrangements [35]. Because the artificial use of the temperature can only expand and contract existing elements, this can not be used to model tissue expansion during time which is a proliferative process. In order to take into account cell division in our model we considered to operate at two different levels, during the first steps of mesh generation or after by updating mesh to allow creation of new daughter cells originating from a mother. Because the second option necessitates dynamical analysis, we only used the first option consisting of mimics cell proliferation (inside a static model) by playing on cell distribution at the first step of organoid generation, creating regions with higher density comparable to higher cell division events(Fig. 3A). This static model, using a probabilistic vision of cell proliferation, was used to simulate shell expansion and deformation over time (Fig. 3B). For that, we used previous active vertex models that have been proposed to study epithelial deformation in multiple biological contexts [36, 37]. These previous studies are generally based on a simple energy functional equation (comparable to a behavior law) to model epithelial monolayer relaxation as described in Farhadifar 2D model:

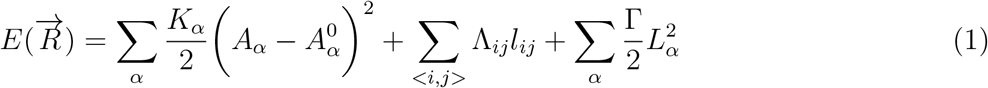

**Figure 3:**
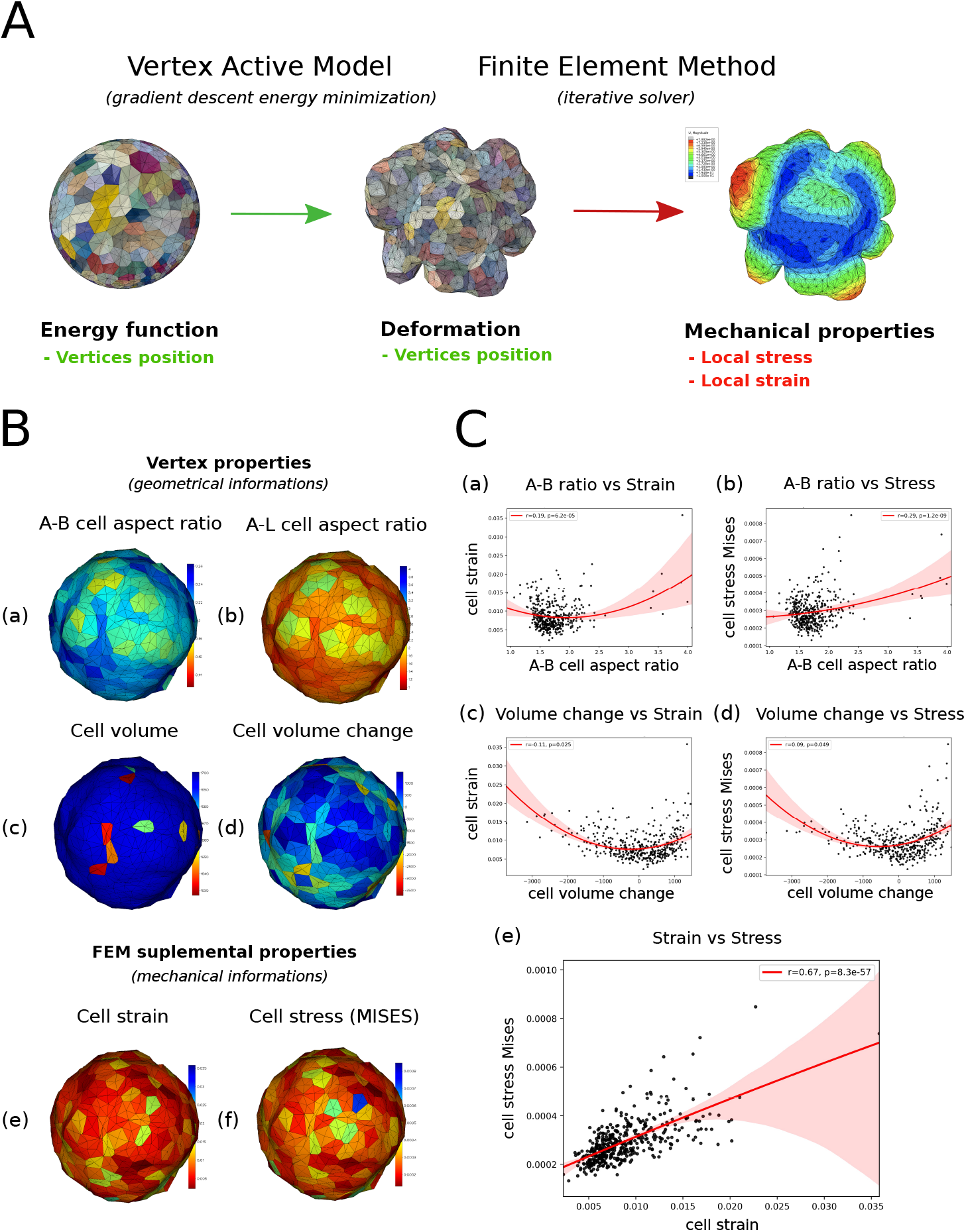
Finite Element Analysis allow to access individual cell mechanical properties in vertex based models. A/ Schematization of the FEM applied to deformations obtained using previously published Vertex Active Model (VAM) resolution. This VAM is a rheologycal model using gradient descent energy minimization to calculate the equilibrium organoid shape assuming the hypothesis that cell volumes are homogeneous inside the tissue [10]). We applied the same displacements obtained with VAM using FEM in order to compute new mechanical properties associated with this shape, cell mechanical stress and strain. B/ Plot of the individual cell properties on 430 cells organoids equilibrated with VAM using parameters presents in the II. Geometrical properties like (a) apical-basal area cell aspect ratio and (b) basal-lateral area cell aspect ratio are directly computed according with the vertices positions that are mesh intrinsic properties. (c) Cell volume (after deformation) and (d) cell volume change (difference between cell volume before and after deformation) are the additional volumes of all volume elements constituting the cell. The use of FEM allow to compute a mean (e) strain value and mean (f) MISES cell stress for each cell of the organoid. C/ Correlative analysis with scatter plot graphical representation comparing (a) apical-basal area cell aspect ratio *versus* cell strain (r=0.19 and p=6.2E-05), (b) apical-basal area cell aspect ratio *versus* MISES cell stress(r=0.29 and p=1.2E-09), (c) volume change *versus* cell strain(r=0.11 and p=0.025), (d) cell volume change *versus* MISES cell stress (r=0.09 and p=0.049) and finally cell strain *versus* cell MISES stress (r=0.67 and p=8.3E-57). Each property is measured and averaged at the cell unit for the 430 cells from the organoid represented in B/. *Red line give the potential non-linear correlation (second order equation) and red area the confidence area for the given values. Correlation coefficients are given using Pearson’s correlation test statistics*.

Where α is cell index, ¡i,j¿ the ensemble of edges, K_α_ the elastic coefficients, A_α_ the areas, Λ the contractility of the L_α_ perimeters and Γ the edge tension [16].

After generalisation and application to a 3D problem [4], this mathematical formulation can evolve to include volume elastic coefficient for cell compressibility, face elastic coefficient and lumen compressibility. The new energy minimum is searched through a gradient descent strategy using the Broyden-Fletcher-Goldfarb-Shanno bound constrained minimization algorithms. This model has evolved during time to be implemented with cell death behavior [38] and cell proliferation as a dynamical process [39]. Here we use the problem formalised in the equation 1 to equilibrate organoid shape by gradient descent energy minimization. Then we solve the system adapting the parameters (II) to fit as best as possible with the organoid shape observed in our matrigel cultures (Fig. 1B, Fig. 4A). Then, we used the result of this vertex model equilibrium to record the displacement of each node and introduce it as a loading displacement for our FEM to run it in parallel (Fig. 3B). By this way, we can compute mechanical properties as Stress and quantitative local Strain efficiently using FEA capabilities. If the two models Vertex anterior model and FEM have a relatively similar geometrical description, constructed around a cell centroids and each piece of the models (faces, edges, nodes) expressed by their affiliation to a specific cell, we can note some differences on the computing part. The integrative level is the edge or the node in the vertex model when the FEM is based on elements we defined in our model, here four different elements with their own proper integration law. This new integrative level is more biologically relevant because we used cellular components (membrane, cytoplasm, cytoskeleton) for the design of our FEM (I).

**Figure 4:**
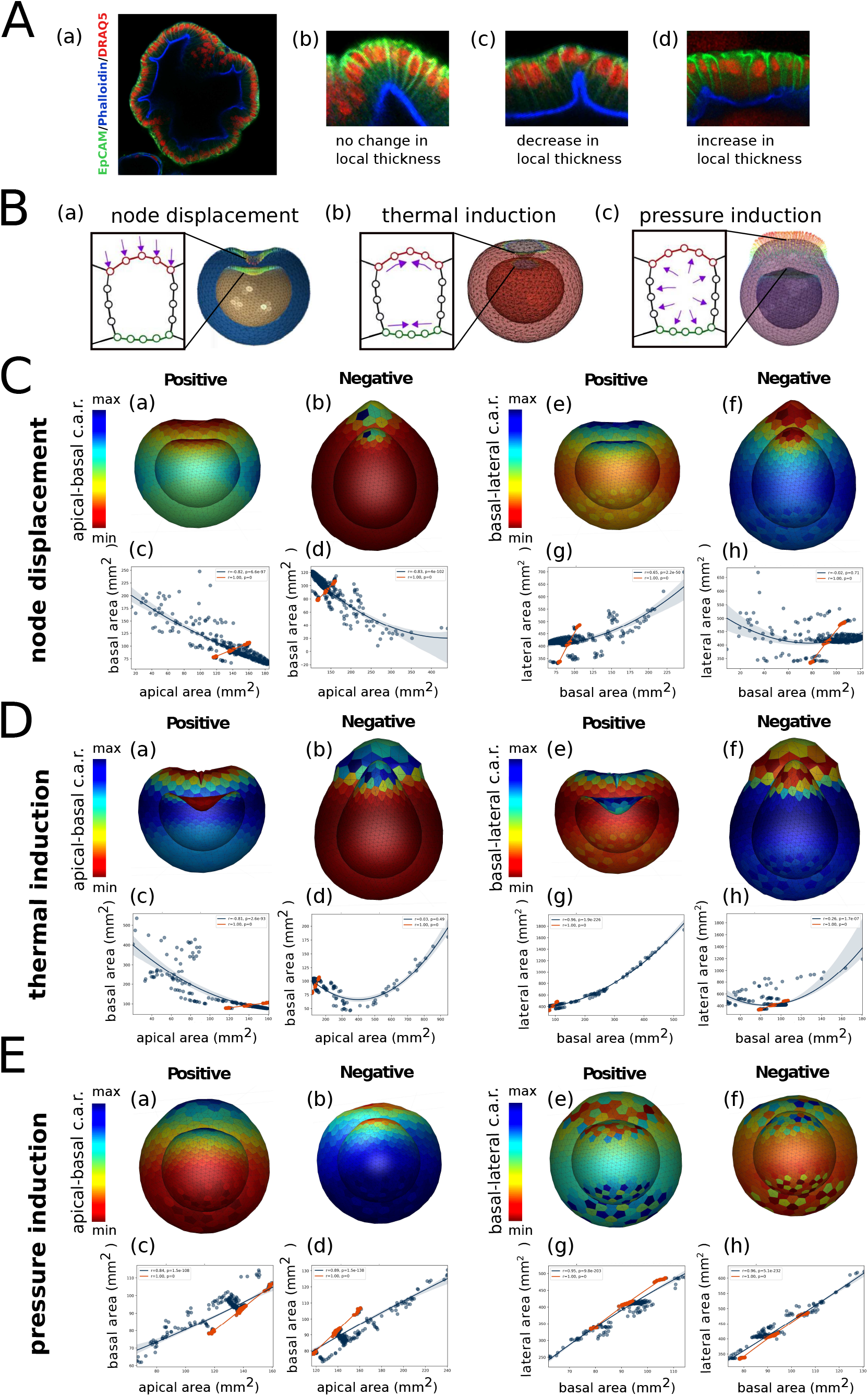
Organoid deformations and cell shape changes links are ensure in FEM. A/ (a) z section of a fixed columnar organoid with phalloidin (blue) immunostaining against actin filaments, EpCAM (green) immunostaining against epithelial cell adhesion molecules and nucleus Draq5 (red) staining. (b-d) Three zoom snapshots realized on the same experiment presenting three distinct cases of deformation with no, negative or positive change in the thickness of the epithelial monolayer. B/ Schema presenting three type of controlled loads. (a) Loads can be apposed as node displacement, here a gradual push on the apical surface resulting in an invagination phenotype. (b) Load can be defined as a specific temperature, contracting both apical and basal surface for negative temperatures, resulting in an invagination phenotype. (c) The last load tested is pressure imposed at the surface of volumic elements, positive pressure dilate the cells and drive an evagination phenotype. C/ Sensibility of the model through controlled node displacement. FEA output deformation of the organoid for (a)(e) positive node displacement (from outside to inside), (b)(f) negative node displacement (from outside to inside) given as (a)(b) apical-basal cell aspect ratio (c.a.r.) or (e)(f) basal-lateral cell aspect ratio (c.a.r.). Scatter distribution of cell geometries for (c)(d) apical-basal c.a.r. (r=0.82, p=6.6E-97 and r=0.83, p=4E-102) or (g)(h) basal-lateral c.a.r. (r=0.65, p=2.2E-50 and r=0.02, p=0.71). D/ Sensibility of the model through controlled thermal induction. FEA output deformation of the organoid for (a)(e) positive thermal induction (negative temperatures), (b)(f) negative thermal induction (positives temperatures) given as (a)(b) apical-basal cell aspect ratio (c.a.r.) or (e)(f) basal-lateral cell aspect ratio (c.a.r.). Scatter distribution of cell geometries for (c)(d) apical-basal c.a.r. (r=0.81, p=2.6E-93 and r=0.03, p=0.49) or (g)(h) basal-lateral c.a.r. (r=0.96, p=1.9E-226 and r=0.26, p=1.7E-7). E/ Sensibility of the model through controlled intra-cellular pressure. FEA output deformation of the organoid for (a)(e) positive intra-cellular pressure induction (increasing intra-cellular pressure), (b)(f) negative intra-cellular pressure induction (decreasing intra-cellular pressure) given as (a)(b) apical-basal cell aspect ratio (c.a.r.) or (e)(f) basal-lateral cell aspect ratio (c.a.r.). Scatter distribution of cell geometries for (c)(d) apical-basal c.a.r. (r=0.84, p=1.5E-108 and r=0.89, p=1.5E-138) or (g)(h) basal-lateral c.a.r. (r=0.95, p=9.8E-203 and r=0.96, p=5.1E-232). *Each property is measured and averaged at the cell unit for the same original 400 cells organoid. Plots show Apico-basal and baso-lateral cell aspect ratio on the apical and basal surface of a side view cutted organoid. Color bar indicate the relative distribution of the cellular aspect ratios (c.a.r.) along the virtual organoid. On correlative distibution plot, blue line give the potential non-linear correlation (second order equation) and blue area the confidence area for the given values and red color is used to indicate the original distribution before deformation. Correlation coefficients are given using Pearson’s correlation test statistics*.

In addition the mechanical relevance of the model is slightly different with the use of global mechanical parameters in the vertex model compared to the possibility to drive each cell with a specific property in the FEM (Fig. 3B). In this conformation, we can observe the regional deformations resulting from the equilibrium of the system and the localisation of cell strain(Fig. 3B). When the only direct exploitable result obtained with the use of the historical vertex model is the resulting shape given by the new position of the nodes, it’s possible with FEA to extract mechanical information in addition to the strain. FEA allows us to check the von Mises [40] yield criterion (MISES), also named equivalent tensile stress, in the post processor that informs us of the resulting stress for each element after deformation. This von Mises constraint is relative to the behavior law we draw in the model, here purely elastic, so can give us the elastic limit. On this deformed organoid, we also explored the possibility to find local geometrical or mechanical descriptors by comparing and performing simple correlatives analysis (Fig. 3C). The two couples of properties presenting a high correlation are apico-basal and apico-lateral cell aspect ratio that are two geometrical descriptors of the cell shape, and MISES cell stress with cell strain. This can point to a relative limitation of the vertex model employed here, where the relationship between stress and strain is relatively linear and independent from the geometry of the cell (Fig. 3D.b). For the following analysis, a part because they are similar in global deformation organoid, we finally exclude apico-lateral cell aspect ratio to previligiate apico-basal cell aspect ratio that can be more informative on cell shape changes and local curvature or deformation of the tissue.

### D. Static model for shell deformation

Because our model is centered on the use of living material, we decided to characterise the link between tissue deformation and cell deformation necessary to design our multi-scale model. We identified three different types of local epithelial shapes present in the advanced columnar organoid (Fig. 4A.a). The first type presents a tissue curvature without important cell shape changes. Here, tissue curvature does not imply any thickness changes (Fig. 4A.b). Second type is more relative to cortical contraction of cells at the apical surface, inducing a thickness decrease at a local point (Fig. 4A.c). Finally, for the third case, an increase of the epithelial monolayer can be observed along with a local cell size increase (Fig. 4A.d). To reproduce these scenarios in our organoid simulation framework, we consider an initial perfectly spherical configuration (Fig. 2A) and induce some perturbations to deform the FEM organoid using Abaqus©, an industrial FEM solver, software capabilities. Our approach consisted in decomposing the shape changes problem into three distinct deformation inputs or loading events that can be complementary or independent (Fig. 4B). They can be resumed as external stress consistent in surface node displacements (Fig. 4B.a), internal stress with the use of pressures exercised on cell surfaces (Fig. 4B.c) or dilatation (contraction) of cortical elements (Fig. 4B.b).

#### a. Cell external forces

Define a displacement at the node level is a common way to compute the resulting strain or constraint with FEM. We used this capability to model epithelial invagination in our FEM organoid. We create a “punch” like function to impose a displacement field on the apical cell surface (Fig. 4B) with an inverted Poisson law distribution in a predefined region, two cell diameter from the initiation. The application of this positive pressure, external to the organoid, conducts to an invagination-like process with a curvature of the basal surface responding non linearly to the deformation of the apical surface (Fig. 4B, Fig. 5A). On the contrary, negative pressure or pulling at the external surface of the organoid conducts to an evagination-like deformation. Curvature analysis has no standard definition and regroups different types of analysis, generally dependent on the domain discretization. This is why we decided to use two different aspect ratio of the cell to quantify the regional deformation of the organoid at the cellular scale as curvature descriptors. Basal-lateral cell aspect ratio is more relevant to survey the evolution of the polarisation (Fig. 1) when apico-basal cell aspect ratio can be more relevant for the local tissue reorganization like local contraction. In this specific case of a punch realized externally to the organoid (Fig. 1B.a), we can see a good correlation between apico-basal cell aspect ratio. Cell strain and Cell stress with a linear correlation between stain and stress correspond to a purely elastic response from this composite material to the loading force imposed. By opposition there is no visible relationship between volume of cells and their deformation. In this case, we can explain that by the fact that deformation occurs essentially by changing the shape of the cell in terms of area more than in volume (Fig. 5A.b).

**Figure 5:**
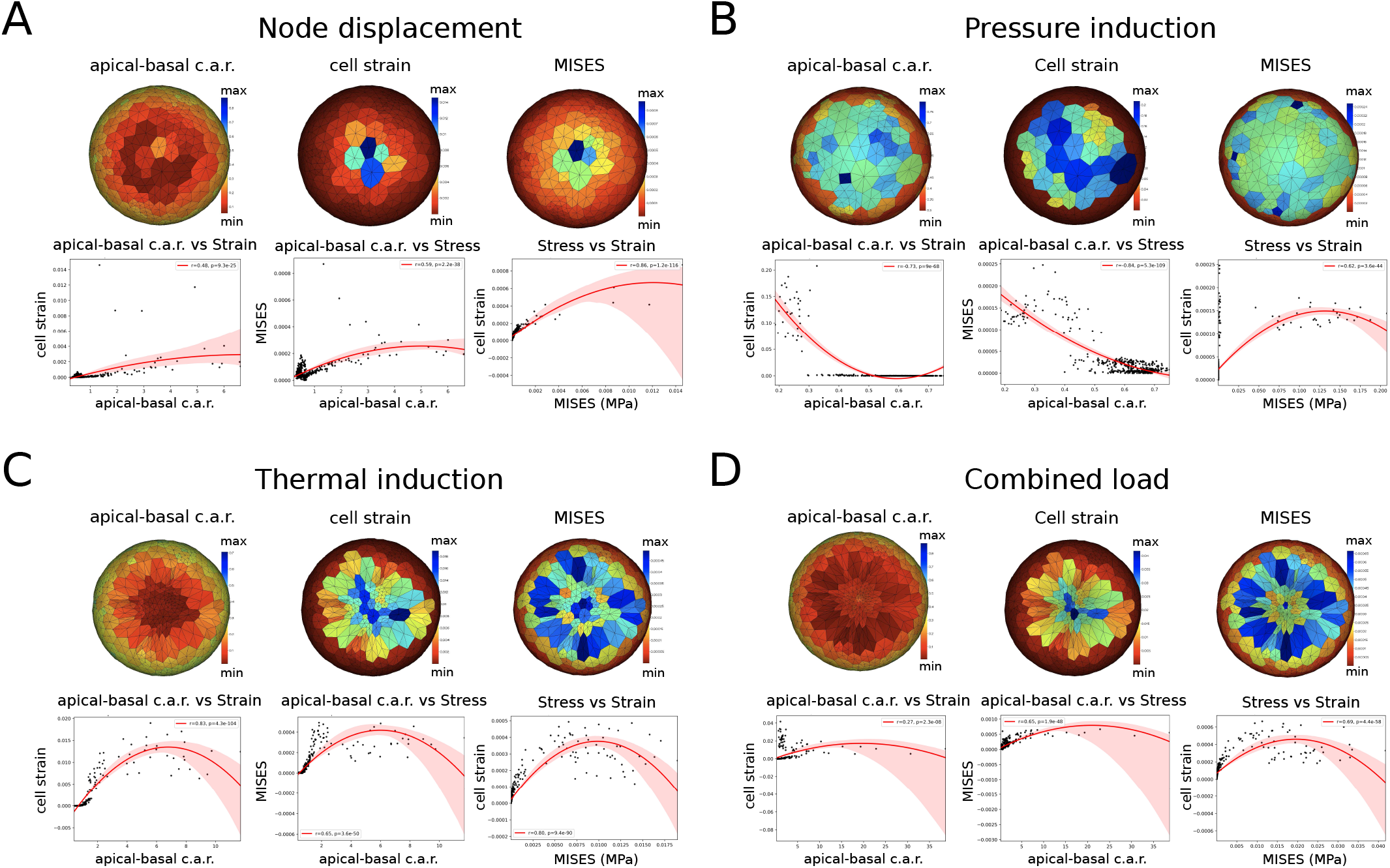
Stress/Strain miscorrelation is dependant of the type of deformation. A/ (a-c) Top view of the deformed organoid following positive ‘node displacement’ induction with plot of the (a) celplot graphical representation comparing (d) apical-basal area cell aspect ratio *versus* cell strain (r=0.48 and p=9.3E-25), (e) apical-basal area cell aspect ratio *versus* MISES cell stress(r=0.59 and p=2.2E-38) and (f) MISES cell stress *versus* cell strain(r=0.86 and p=1.2E-116). B/ (a-c) Top view of the deformed organoid following positive ‘thermal induction’ with plot of the (a) cell strain properties, (b) cell MISES stress and (c) apico-basal cell aspect ration. Correlative analysis with scatter plot graphical representation comparing (d) apical-basal area cell aspect ratio *versus* cell strain (r=0.73 and p=9E-68), (e) apical-basal area cell aspect ratio *versus* MISES cell stress(r=0.84 and p=5.3E-109) and (f) MISES cell stress *versus* cell strain(r=0.62 and p=3.6E-44). C/ (a-c) Top view of the deformed organoid following positive ‘intra-cellular pressure’ induction with plot of the (a) cell strain properties, (b) cell MISES stress and (c) apico-basal cell aspect ration. Correlative analysis with scatter plot graphical representation comparing (d) apical-basal area cell aspect ratio *versus* cell strain (r=0.83 and p=4.3E-104), (e) apical-basal area cell aspect ratio *versus* MISES cell stress(r=0.65 and p=3.6E-50) and (f) MISES cell stress *versus* cell strain(r=0.80 and p=9.4E-90). D/ (a-c) Top view of the deformed organoid following ‘combined’ induction, an additive combination of the three precedent inductions, with plot of the (a) cell strain properties, (b) MISES cell stress and (c) apico-basal cell aspect ration. Correlative analysis with scatter plot graphical representation comparing (d) apical-basal area cell aspect ratio *versus* cell strain (r=0.27 and p=2.3E-8), (e) apical-basal area cell aspect ratio *versus* MISES cell stress(r=0.65 and p=1.9E-48) and (f) MISES cell stress *versus* cell strain(r=0.69 and p=4.4E-58). *Each property is measured and averaged at the cell unit for the same original 400 cells organoid. Plots show specified properties on the top view organoid, where induction occurs. Color bar indicate the relative distribution of properties along the virtual organoid. On correlative distribution plot, red line give the potential non-linear correlation (second order equation) and red area the confidence area for the given values. Correlation coefficients are given using Pearson’s correlation test statistics*.

#### b. Thermal model of cell contraction

When forces at apical surface of the cell are well characterised for apical contraction requirement during invagination of the epithelium, basal surface of a cell is more commonly associated with cell expansion where basal acto-myosin network is actively disassembled. This could be a way in which basal relaxation could regulate or be necessary for subsequent apical constriction during invagination [41]. We simulated differential cortical contraction at the apical and basal surface of the cells defining a specific expansion rate and temperature to reach during the simulation for cortex elements I. We realized induction with an extremely low temperature to induce (Fig. 4C.b) invagination like deformation of the organoid and the opposite high temperature induction to expand cortexs and create evagination (Fig. 4C.b’). We can observe a drastic apico-basal cell aspect ratio change in the induced area and a loss of the linear correlation between this two areas relationship, suggesting important individual cell shapes changes. If stress (von Mises constraints) accumulated by the cells is not correlated with apico-basal cell aspect ratio or cell strain, volume is surprisingly correlating with all other properties (Fig. 5B). This can be explained by the fact that inducing both apical and basal contraction or expansion, even if differential mechanical properties (expansion rate five times more in apical than in basal) is sufficient to create direct volume expansion of the cells. We can imagine adding lateral contraction or increasing the differential contraction to be more relevant biologically but his results suggest that cell contraction can be modelled by thermal expansion and conduct cell rearrangement and changes in cell shape that mimic living cells behaviors [42].

#### c. Cell internal pressures

Osmotic forces are another important component emerging from mechanical models [43]. If forces at the apical surface of the cell are well characterised as apical contraction requirement during invagination of the epithelial shell, surprisingly, control of the hydrostatic pressure is much less studied in the context of tissue deformation. In our model, we considered cells as more or less purely elastic but we do not constraint the compressibility if it’s not through the Poisson ratio attribute in complement to the Young’s modulus. However, we can change this osmotic pressure using pressure forces directly on specific cell surfaces as loading parameters in our FEM (Fig. 4B.c). We decided to try this by exerting a negative pressure (directed against the external surface of the cells, Fig. 4C.c), or a positive pressure (directed against the internal surface of the cells, (Fig. 4C.c’), to respectively contract or expand the volume of the cells on a localized region of our simulated organoids. This two perturbation results in a global change of the induced cells with a shift in the apico-basal cell aspect ratio, with a similar decrease of apical and basal cell area for negative induction and an inverse proportional increase when inner cell pressure increases. The model meets expectations concerning volume changes (Fig. 4C). If we analyze the correlation between geometrical and mechanical cells properties we can see that there is a correlation between volume and apico-basal cell aspect ratio as seen in previous analyses. Increase in volume correlates with an increase of both apical and basal area. But a more interesting observation is the presence of not one but three regimes of correlation between cell volumes and strain. Indeed it can be understood as three different cell behaviors depending on the localisation of cells inside the induced area. At the center, induced cells that are in contact with cells that are also induced. And two populations at the periphery of the induced area, first induced cells inside the border contacting not induced cells and to finish cells outside the induction area sharing interfaces with induced cells(Fig. 5C).

With all this observation we can conclude that apico-basal cell area ratio is a good descriptor for local deformation when volume changes have a more restricted relationship relative to osmotic changes.

## III. DISCUSSION

Personalized medicine consists in precise diagnosis and the application of specific treatments to each patient. Organoid cultures represent a perfect opportunity to assess patient specificity, offering a lot of biological materials from a minimum of biopsy explant. Organoids further surpass 2D cell cultures in terms of structural resemblance, useful in the case of biomechanical studies [44]. Another advantage of organoids is to provide faster and more robust outcomes which are more readily accessible for live imaging. This is why image-based screens on organoid cultures have emerged as an effective and accurate model system to study pathogenesis or to propose personalized diagnosis and treatments[45].

Research into biomechanics has not only provided new insights for a better elucidation of the mechanisms behind disease progression, clarifying the differences between diseased and healthy cells, but has also provided important knowledge in the fight against these diseases. However, there is a lack of information on the relationships between architecture, mechanics and biological functions, especially in organoids cultures. Moreover, measuring sub-cellular mechanical properties require invasive experiments like atomic force microscopy (AFM) [46] or traction force microscopy (TFM) [47] that are not fully compatible with organoids 3D architectures and culture conditions.

For all these reasons, we decided to develop a new virtual model of organoid integrating geometrical and mechanical properties at the cellular level, associated with structural FEA to access the physical behaviour of a living epithelial tissue.

Taking advantage of exciting opportunities offered by organoids, we decided to focus our attention on imaging the precise tissue architecture at a cellular level. Numerous 3D in *silico* models of epithelial tissue or organoids use vertex model simulations with generic formulation of energies. If these models can be effectives to describe some biological morphogenetic processes, they suffer from limitations in terms of cell shape. Usually, in a vertex mechanical model with individual cell shape description, analytical predictions are difficult to obtain because of the exponential number of parameters depending on the total cell number. FEA represents a better alternative to solve analytical predictions in the case of complex tissues like large columnar organoids. Associated with the 3D vertex model, we can compute mechanics from the geometry and create a 3D object onto which quantitative geometric or mechanical data can be projected. All the individual subcellular properties can be used to drive unsupervised analysis in order to fill the gap between cell shape and cell mechanics.

In the study of structural architecture, one of our main proposals is that epithelial tissue mechanical information is recapitulated in tissue and cell shape. To ensure correct computing performance in terms of solving time, without compromising cell geometry resolution, we choose to use a model designed around the minimum of sub-cellular elements to construct our virtual tissue model. This virtual cell construction involved cell components, namely, cytoplasm, membrane and cortex, each associated with a specific element formulation and individual material properties. As a result of our simulations, we can observe that our simple model architecture, first based on vertex and then translated into finite elements, is not only compatible with precedent designed models but additionally provides mechanical information regarding cell or tissue constraints, deformation and materials properties. One strong limitation of the general formulation proposed in the actual vertex model lies in the fact that geometry is frequently imposed on cells. This is not an issue for tissue composed of regular cell shapes like in Drosophila [48], but this can be less representative of the cell variability present in organoids and creates artificial behavior decorrelated from the geometry (figure 3C).

Another important result can be the validation of apical-basal cell area aspect ratio as a simple descriptor to survey local curvature in organoids deformations. Basal-lateral cell area aspect ratio is more relevant to survey the temporal evolution of the organoid and to determine the transition from cystic to columnar organoids.

We also note that change in volume of the cells (figure 4D) can be a good reporter of the osmotic changes and can have a real impact on the tissue thickness associate with changes in local curvature.

There is a potential interest to play on the volume, when new born cells have to grow after cell division. In the special case of cell division, cell shape variation can be drastically dynamic during mitosis compared to the interphase portion of the cell cycle. During mitotic cell rounding, cells abandon their elongated shape and contract into a spherical morphology in combination of apical cell-cell junction maintenance and the increase of the surface tension of the actin cortex [49, 50]. The use of our quadratic elements allows more freedom in the cell deformability and the possibility to obtain round cells, after refinement by increasing the number of elements per cell. However this refinement has a dark side effect increasing drastically the number of integrating points for the computation and so the computation time.

We additionally observed that we can modulate tissue thickness, without changing lateral membrane with the contribution of differential apical-basal contractility or internal (osmotic) cell pressure when an important contribution of the lateral membrane contractility was observed in other studies [51]. If we can explore the cell specific shape more precisely along the apical-basal axis and perform accurate measurements of the height profile, necessary to corroborate the direct link between apico-basal ratio and strain accumulate by the cells, importance of the lateral contractility seems to be more interesting to explore in a more complex model, with individual cells and contact modelled between each cells. However, surveying the evolution of this shape is a challenging task, given the high aspect ratio (thickness/diameter) of the cells during polarisation and when tissue is in a really compact, budding phase.

Our numerical results indicated that apical purse-string tension, when applied together with volume preservation, induced a reduction of the height, due to the expansion of the tissue. We point out that in our model, lateral contractility is homogeneous along the entire tissue. This is why we do not modulate this parameter. Due to the apico-basal polarisation of the cells and the evolution of this polarisation during organoid development (figure 1), the lateral contractility may not be homogeneous along the thickness. Further discretization of the monolayer along the apico-basal axis seems necessary to simulate the specific localization of lateral myosin on the apical side. Moreover, myosin phosphorylation and ectopic myosin activity, could promote contraction and detachment of newly formed interfaces resulting from the T1 or T3 transitions [52].

Interestingly, it has also also shown that tissue tension and contractility are also driven by the tissue fluidity, the intercalation of cells during time [53]. This points to the necessity to push more flexibility in cell geometry, and our model goes in this way, for dynamical analysis. The analysis of specific molecular components and the evolution of the material properties after cell rearrangement including T1,T3 transitions, cell division and cell polarisation is left for further investigations.

The second aspect of our model is the use of FEM to acquire physical information based on the geometry. We explored the FEA capabilities using calibrated loading cases translated in observable deformations and from which we can compute some mechanical properties. We propose to reduce the dazzling variety of local tissue shape or local deformation, coming from the assumption that the shape originates from a standard spherical conformation, using only three scenarios. This simple clustering case defined using thickness changes is sufficient to characterize all the observable deformation. In the case of image analysis, we can obtain this displacement from live imaging registration or directly use finite element analysis updated (FEMU), dealing with time [54]. The remaining unknown parameters are forces involved and materials parameters. Following this observation, we proposed to consider three types of highly parameterizable loading cases. If these loads are not considered as an exhaustive list of possible perturbation, she allows by a combination of these different input to create a large collection of phenotypes sufficient to create the conditions of the three scenarios described. Simulating the recorded deformation using our simple loading cases can be used to approximate the relative distribution of the physical parameters and determine if cells are more or less elastic inside the tissue. We can easily use that for future unsupervised study in order to define the area of interest in the imaged organoids. All together these results confirm the flexibility of our FEM formulation to create complex deformation. This was the case with previous vertex models allowing to create high deformations for morphogenetic events [4, 55] or artificial tissues [14, 39, 56] without using FEM analytic power. FEM interest in biology is not new, FEM was previously used to model morphomecanic at a meso-scale level [57] and FEM was recently implemented on a nice platform for quantifying plant morphogenesis in 2.5D, MorphoGraphX, that can be used for image analysis, and using surface elements [58]. Our study makes the link between these two approaches, simulation of a relevant 3D tissue architecture at the sub-cellular level and efficient mechanical analysis. We expect now that more accurate experimental measurements and model enhancements will allow us to quantify cell mechanical contributions and regulations more closely, in order to enhance our virtual model.

Our virtual proposition can be seen as a proof of concept allowing us a nice definition and validation of the FEM model of organoid. Future investigation will be focused on the usage of this wonderful tool to perform inferring analysis using inverse formulation of the problem linking cell shape and mechanical parameters [59–61].

In conclusion, our multiscale model, from subcellular elements to full-size organoid proposes a new way to model epithelial tissue that considers cell mechanics using cell based FEM. In this model, mechanical parameters are based on material law description like in engineering study. This approach is particularly adapted for inverse problems allowing to infer forces or material properties from the geometry with efficient computing and mathematical validates tools. We show that combined simple cell geometric descriptors, simple mechanical descriptors like cell stress and cell strain and controlled forces allow us to model the local deformation of the tissue. Our design of a complex and global problem of morphogenesis, tissue folding, adapted to organoid structure is a first step for an image based physical inference framework and could help future direct image analysis and machine learning implementation diagnosis in personalised medicine. Indeed, the simplified geometric formulation of the model makes it compatible with a huge range of image solution as long as we can reconstruct the cell shape.

Globally, these numerical and geometrical simulations would be necessary to investigate the role of cell singular or local environment mechanics changes in pathophysiology studies.

## IV. METHODS

### A. Human samples

Biological samples were obtained from patients at the Toulouse University Hospital. Patients gave informed consent and were included in the registered BioDIGE protocol, approved by the national ethics committee (NCT02874365) and financially supported by the Toulouse University Hospital. Colonic samples were obtained from biopsies of patients undergoing endoscopy.

### B. Organoid culture and analysis

Colorectal crypt isolation was performed as described previously [62]. Fresh Matrigel (Corning, 356255) was added to isolated crypts. 25μL of Matrigel containing 50 crypts were plated in each well of a pre-warmed 8-well chamber (Ibidi, 80841). Once the Matrigel had polymerized for 20 min at 37°C, 250μL of culture medium was added to each well as described previously [62]. Cultures were followed over two weeks for analysis. Regarding immunofluorescence staining, organoids were fixed with a PBS solution containing 3,7% of formaldehyde solution for 5 min at 37°C. Then, organoids were washed in PBS and permeabilized with a PBS solution containing 0.5% Triton X-100 for 20 min at 37°C. After washes, DNA was stained with Draq5 (Ozyme, 424101, 2 μM) for 30 min, membranes were stained using anti-EpCAM antibody (Cell Signaling Technology, VU-1D9) and actin network using Alexa Fluor 594 phalloidin (ThermoFisher, A12381, 40nM). Finally, cover glasses with Matrigel domes were immersed in PBS for imaging. For live experiments, DNA was stained with the Spy650-DNA probe (Spirochrome, SC501). Images acquisition were performed using a two-photon microscope (Bruker 2P+, 20X diving lens objective) for fixed experiments and with the Opera Phenix HCS microscope (Perkin Elmer, 40x objective) for live experiments. Images were analysed using the ImageJ software from FiJi [63].

### C. Model Implementation

We used python libraries numpy, scipy to generate the mesh, this mesh is store in Dataset (panda) format and can be export directly in hdf5, obj (for surfaces) or vtk format using meshio package. A large part of the python codes are using Tyssue library for monolayer creation [64], .Cgal (via python wrapper) and pygalmesh are used as cell/cell contact, remeshing and cell division solution and VTK for the visualisation. A conversion to .inp (input file) Abaqus© was done in python to create our definitive FEM virtual organoid. Readout of the Abaqus© FEA was performed in python with the development of a .fil reader and the use of vedo for graphical representation [65].

## Author contributions

AF and FBu designed the project. JL, FBu, AF and SS contributed to the conception of the original idea. JL developed the theory and the finite element model, performed simulations, analysed data sets and results, and wrote the manuscript; EM coordinates the BioDIGE protocol, screened the patients, performed the endoscopy and biopsies; BioDIGE is supported by Toulouse University Hospital. DH established the organoid cultures; DM performed organoid cultures, immunofluorescence stainings, images acquisition and images analyses; AF developped the theory and its connection to the organoid model; AF, SS and FBu developed the theory, did the interpretation of the results and wrote the manuscript. All authors contributed to the discussion and interpretation of the main results.

## Acknowledgments

This collaborative work was funded by two Cancer Plan projects: Mocassin - Biosys 2017, granted to AF and Melchior - Mic 2020, granted to FBu. JL is supported by both, DM is supported by Plan cancer Melchior-MIC 2020. They also wish to thank the patients who agree in giving their tissue for research purpose and allowed this study.

